# Collapse of the hepatic gene regulatory network in the absence of FoxA factors

**DOI:** 10.1101/2020.03.31.016006

**Authors:** Yitzhak Reizel, Ashleigh Morgan, Long Gao, Yemin Lan, Elisabetta Manduchi, Eric L. Waite, Amber W. Wang, Andrew Wells, Klaus H. Kaestner

**Affiliations:** Department of Genetics, University of Pennsylvania, Philadelphia, PA; Epigenetics Institute, University of Pennsylvania, Philadelphia, PA; Department of Biostatistics, Epidemiology & Informatics, University of Pennsylvania, Philadelphia, PA; Department of Pathology and Laboratory Medicine, University of Pennsylvania, Philadelphia, PA

## Abstract

The FoxA transcription factors are critical for liver development through their pioneering activity, which initiates a highly complex regulatory network thought to become progressively resistant to the loss of any individual hepatic transcription factor via mutual redundancy. To investigate the dispensability of FoxA factors for maintaining this regulatory network, we ablated all FoxA genes in the adult mouse liver. Remarkably, loss of FoxA caused rapid hepatocyte dedifferentiation manifested by a massive reduction in the expression of key liver genes. Interestingly, expression of these genes was reduced back to the low levels of the fetal prehepatic endoderm stage, leading to necrosis and lethality within days. Mechanistically, we found FoxA proteins to be required for maintaining enhancer activity, chromatin accessibility, nucleosome positioning and binding by HNF4α. Thus, the FoxA factors act continuously, guarding hepatic enhancer activity throughout life.

## Introduction

Developmental programs are composed of multiple successive signals that bring about accurate and robust patterns of gene expression necessary for organ formation. The role of pioneer factors is to initiate transcriptional activity through nucleosome re-positioning at *cis*-regulatory sequences (Zaret and Mango 2016). Nucleosome repositioning is essential in order to allow the recruitment of the full complement of cell type-specific transcription factors to a specific set of enhancers (Zaret and Mango 2016). Once transcriptional activation is achieved, pioneer factors are not considered necessary for the maintenance of transcriptional execution, either because they are replaced by other transcription factors or because these regulatory sequences are fully activated (Spitz and Furlong 2012). There are only a few pioneer factors that are explicitly known as competency factors, required only for activation during early embryonic development and not at later stages (Liang et al. 2008). An example is Zelda, which in *Drosophila melanogaster* is required for enhancer activation only in early embryos, but replaced by other transcription factors occupying the same regulatory sequences later in life (Liang et al. 2008).

To date, it is not clear if exclusively competency factors such as Zelda exist in vertebrates, though the FoxA winged helix transcription factors have been suggested to play such a role (Lee et al. 2005; Jacobs et al. 2018). FoxA proteins, previously defined as “paradigm pioneer factors” (Lee et al. 2005; Li et al. 2012; Iwafuchi-Doi et al. 2016), are critical for the induction of the liver primordium from foregut endoderm (Lee et al. 2005), can bind to nucleosomal DNA both *in vitro* and *in vivo*, occupy *cis*-regulatory sequences before activation of the associated genes, and recruit ATP-dependent chromatin remodelers (Li et al. 2012; Iwafuchi-Doi et al. 2016). Circumstantial evidence suggests that FoxA proteins, similarly to Zelda, are not essential for maintaining the gene network of terminally differentiated cells. First, after FoxA executes its pioneering activity at the prehepatoblast stage, progressively more transcription factors are activated and often bind to the same enhancers already engaged by FoxA proteins, creating a highly complex network thought to be resistant to loss of binding by any specific hepatic transcription factor (Kyrmizi 2006). Indeed, while hepatic depletion of HNF4α during embryonic development results in lethality, ablation of the same transcription factor during adulthood does not cause death (Kyrmizi 2006; Bonzo et al. 2012). A similar phenomenon was observed for C/EBPα, for which only its early ablation causes lethality (Inoue et al. 2004). Second, while FoxA1/A2 are critical for the formation of the liver primordium (Lee et al. 2005) and embryonic survival, their removal at later stages of development has a much milder phenotype which includes cholestasis under specific conditions (Li et al. 2009). Third, recently Thakur and colleagues analyzed the epigenetic role of HNF4α and FoxA2 in adult liver and concluded that FoxA2 has no significant role in maintaining enhancer activity in the adult liver (Thakur et al. 2019). Thus, the view in the field is that FoxA proteins main function is to act mainly as competency factors to establish a stable enhancer network during embryonic development. In this study, we evaluated this concept by deleting all FoxA proteins in the adult mouse liver and assaying network stability.

## Results

### FoxA proteins are required to maintain liver homeostasis

We depleted all three FoxA genes in 8 week-old mice by injecting adeno-associated virus 8 (AAV8) carrying the gene for Cre recombinase under the control of the hepatocyte-specific thyroid-binding globulin (*Tbg*) promoter to adult FoxA1^L/L^/FoxA2 ^L/L^/FoxA3^−/−^ mice (Fig. 1A, ‘FoxA triple null’ below for short) and validated FoxA depletion (Supplemental Fig. 1A). As controls we injected the same transgenes with AAV8 expressing GFP. FoxA triple null mice started losing weight 15 days after virus injection. 20% of these mice died within the next four days and the remainder had to be sacrificed following ethical guidance (Fig. 1B). Livers were cirrhotic (Fig. 1C) and plasma levels of the hepatocyte enzymes alanine aminotransferase and alkaline phosphatase were dramatically elevated, indicating severe liver injury (Fig. 1D). Acute midzonal coagulative necrosis was present in individual or clustered hepatocytes as well as multifocal hepatocellular swelling and clearing (Fig. 1E, and Supplemental Fig. 1B) indicating internal liver failure. In addition, we observed multifocal random mixed lymphocyte infiltration, demonstrating that inflammation followed cell destruction and not the opposite. Interestingly, in liver sections of FoxA triple null mice taken shortly after AAV-Cre injection, there was only mild multifocal hepatocellular atrophy with centrilobular hepatic cord hypercellularity, corresponding to a higher Ki67 labeling index at this stage (Fig. 1E and Supplemental Fig. 1B and 2). Overall, the striking phenotype following FoxA depletion in the mature liver demonstrates its necessity for liver homeostasis, showing that FoxA’s critical role is not restricted to early organ development.

**Figure 1.**
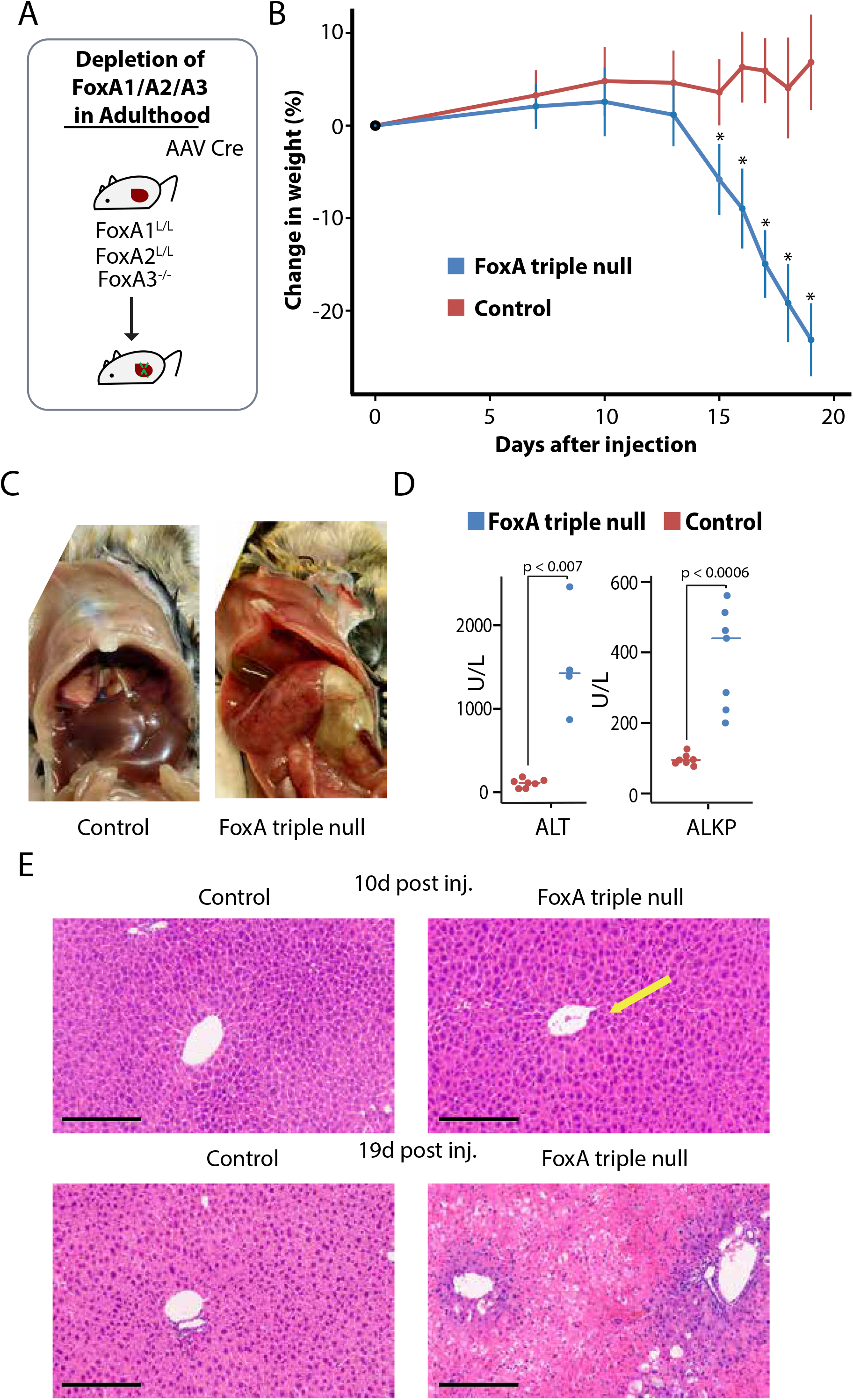
Mice deficient for FoxA proteins succumb to liver failure. A. Description of gene ablation model. B. Rapid weight loss after acute FoxA deletion (* p<0.0005 Wilcoxon-Mann-Whitney test, n≥=8). C. Liver anatomy of FoxA triple nulls and controls. D. Blood levels of ALT, ALKP (p<0.01 for all comparisons, Wilcoxon-Mann-Whitney test, n≥4). E. Liver histology demonstrates a mild phenotype 10 days after AAV-Cre injection in FoxA triple nulls compared with controls, but massive necrosis 19 days following injection. Scale bar = 200 μm.

### FoxA ablation in adult mice eliminates expression of key liver genes

Next, we compared hepatocyte gene expression levels of FoxA triple null livers to controls or to models of limited FoxA deficiency established previously (Li et al. 2009). We performed this analysis seven days after AAV injection before hepatic necrosis had begun, to capture only the most relevant and direct FoxA targets. We discovered 663 genes specifically downregulated in the FoxA triple null livers but not in partially FoxA depleted models (Supplemental Fig. 3A). Gene ontology analysis identified key liver processes associated with the downregulated genes such as lipid metabolism, acute phase response, coagulation and complement system, metabolism of xenobiotics, bile acid synthesis, and the urea cycle (Fig. 2A). Many key liver genes like albumin and afamin (*Afm*) displayed reduced expression in the FoxA deficient liver (Fig. 2B and Supplemental 3B), and a subset of these showed near total loss of steady state mRNA, with decreases of more than 30-fold (Fig. 2B). Among these was the transport protein transthyretin (*Ttr*) that carries thyroid hormone and retinol. The Ttr promoter contains the FoxA binding site that was used to purify the FoxA proteins (then termed ‘hepatocyte nuclear factor 3’) more than 30-years ago (Costa et al. 1988), but only now shown to be dependent on FoxA for gene activation (Fig. 2B). Among the FoxA-controlled-genes, Arginase 1 (*Arg1*) is most relevant to the pathophysiology of the mutant mice. *Arg*1 catalyzes the last step of the urea cycle and is required for the hydrolysis of arginine to ornithine and urea, and its expression is reduced by 40-fold in FoxA mutant mice (Fig. 2B). Complete liver-specific deficiency for *Arg1* is lethal due to hyperammonemia and liver damage (Sin et al. 2013). Since we observed a dramatic reduction of *Arg1* expression and a hyperargininemia, hypouremia and hyperammonemia similar to those present in *Arg1* null mice, we attribute the lethality of the FoxA triple null mice to this defect (Fig. 2C and Supplemental Fig. 4D). We found additional key metabolic genes with massively reduced expression such as *Pemt* that encodes an enzyme which converts phosphatidylethanolamine to phosphatidylcholine, *Apoa1* which is a major component of HDL particles in the plasma, and Cyp8b1 that controls the balance between cholic acid and chenodeoxycholic acid (Fig. 2B). The fact that additional key metabolic genes controlling cholesterol homeostasis are also strongly FoxA dependent is reflected in significantly altered lipid and carnitine levels, which likely contribute to the severe liver damage in FoxA deficient mice (Supplemental Fig. S4). Interestingly, we found that the midzone of the hepatic lobule, i.e. the area with the most significant pathology (Fig. 1E and Supplemental 1B), exhibits highest expression levels of genes completely dependent of FoxA, including *Arg1, Pemt, Cyp8b1* and *Apoa1*, as determined by zonation gene expression profiling (Figure 2D, zonal expression data was adapted from (Halpern et al. 2017)).

**Figure 2.**
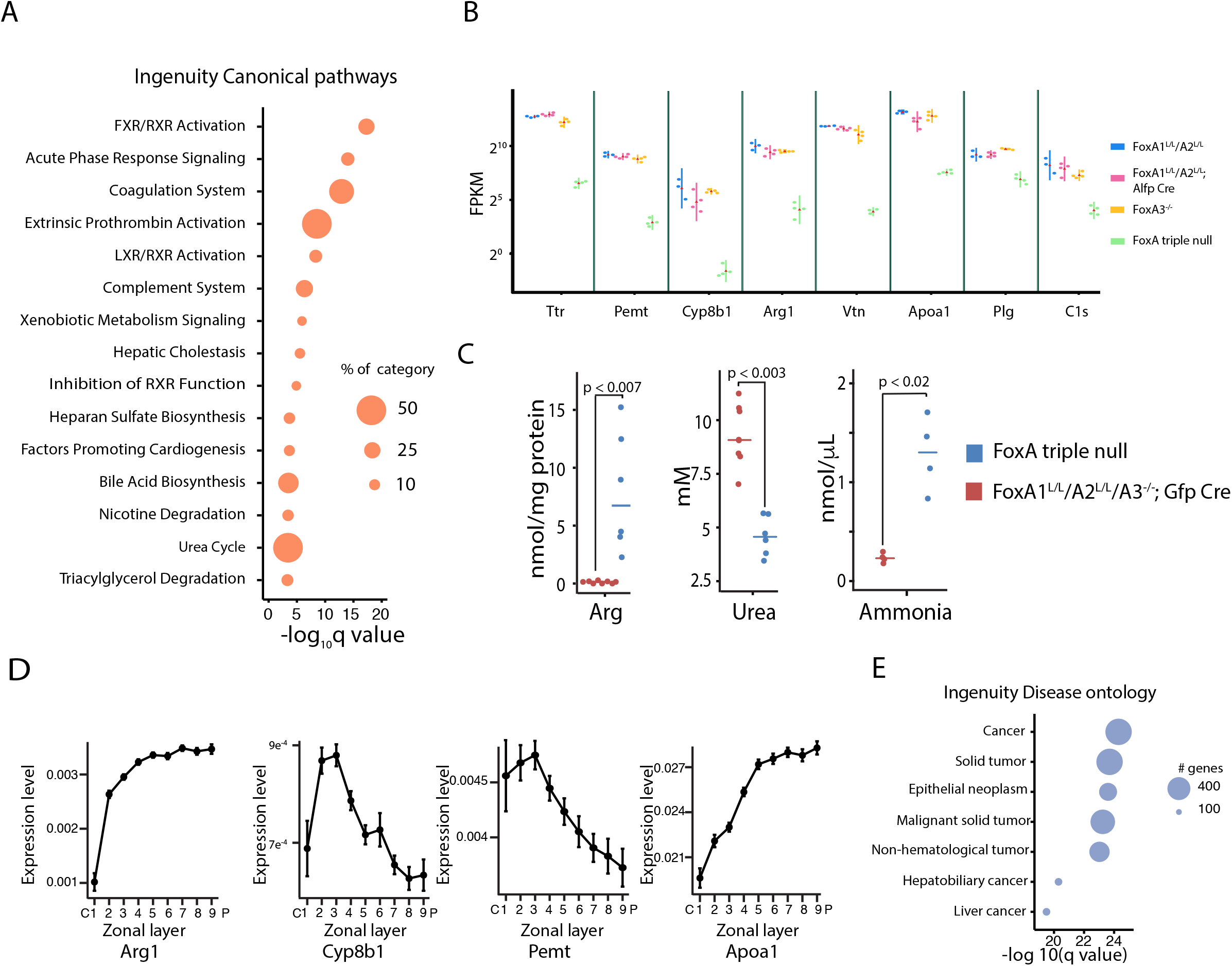
Key liver genes are dependent on FoxA proteins in the adult liver. A. Canonical pathways associated with down-regulated genes in the FoxA triple null. B. Examples of key hepatic genes down regulated in the FoxA triple null liver. Note the log scale (green triangle is the mean, adjusted pvalue < 10^−40^ for all comparisons between FoxA partial depletion or controls and FoxA triple null). C. Hepatic levels of Arginine and Urea and blood levels of Ammonia (Wilcoxon-Mann-Whitney test, n≥ **4**). D. Zonal distribution of four key genes down-regulated in the FoxA triple-null showing high expression in the midzonal region (Arg1 and Cyp8b1 were reprinted with permission from Halpern et al. 2017). E. Disease ontology of upregulated and down regulated genes using ingenuity.

We also found 513 genes to be upregulated in FoxA triple null livers (Supplemental Fig. 3C). Interestingly, gene ontology analysis of the upregulated genes indicated that many of these are associated with cancer and related proliferative pathways (Fig. 2E). This gene signature explains the hepatic hypercellularity and increased proliferation rate immediately following AAV-Cre administration reported above (Fig. 1E and Supplemental Fig. 2) and reflects the attempt of the injured liver to restore functional tissue mass.

### FoxA proteins maintain the hepatic differentiation state

Since FoxA proteins are central to the induction of gene expression during early pre-hepatic endoderm development (Lee et al. 2005), we wondered how many of the genes down-regulated in the triple null model are developmentally induced. Strikingly, we found that all genes with more than a 30-fold decrease in the mutants (Fig. 2B) are massively activated during liver development, and in most cases, their expression levels regress back to those of the pre-hepatic endodermal stage (Fig. 3A, gene expression of endodermal and hepatoblast were obtained from (Nicetto et al. 2019) and early postnal hepatocytes form (Reizel et al. 2018)). When we extended this analysis to all 663 significantly downregulated genes, we found that 63% of these are developmentally induced and 24% are deactivated in the triple null mutants back to fetal levels (Fig. 3B-C), demonstrating a clear dedifferentiation phenotype. This is unlike the genes that are upregulated in the FoxA triple null livers, of which 77% are not developmentally controlled and whose activation is associated with transcription factors involved in the injury response (Supplemental Fig. 5A-D). Overall, our transcriptome analysis demonstrates that the FoxA proteins are continuously required to maintain the differentiated state of hepatocytes by sustaining the expression of crucial liver genes and by blocking the expression of early developmental genes (Supplemental Fig. 5C-D).

**Figure 3.**
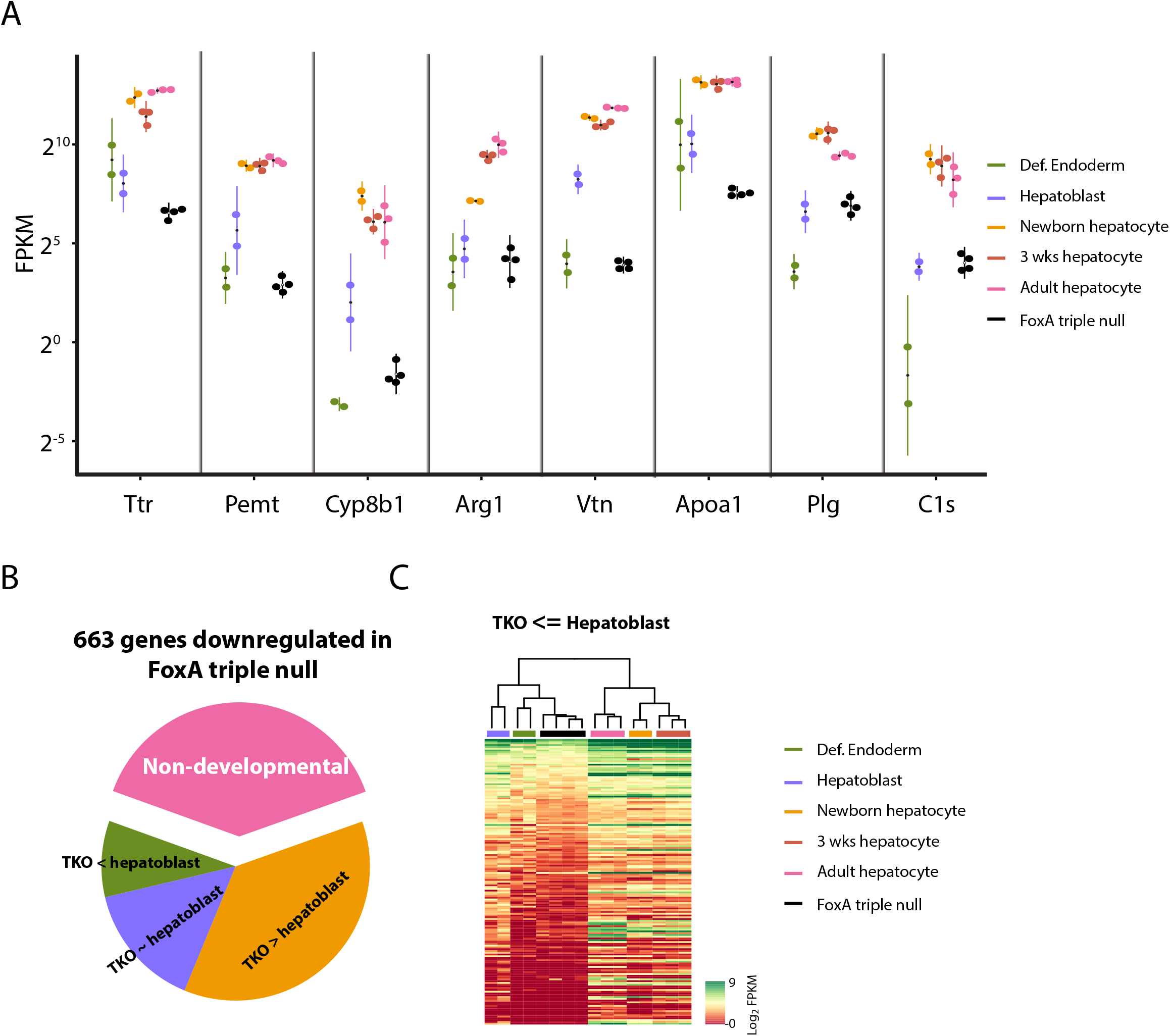
Adult ablation of FoxA factors reduces expression of key liver genes back to early developmental stages. A. Expression of FoxA target genes is reduced back to early fetal stages following acute FoxA ablation (def. = definitive). All genes are statistically significant lower in FoxA triple null compared with hepatoblasts (adjusted p value < 0.001) except for Arg1, Plg and C1s. B. The majority of FoxA dependent genes in the adult liver are induced during liver development, and many are deactivated in the FoxA triple null liver back to total hepatoblast levels (adjusted p value < 0.01, details Materials and Methods). C. Heatmap demonstrating downregulated genes regressing back to hepatic or prehepatic stage in the FoxA triple nulls.

### FoxA binding sites are associated with downregulated genes

In order to find association between FoxA binding sites and downregulated genes in FoxA triple null, we focused on FoxA3 sites in liver deficient for FoxA1 and FoxA2 (for explanation, Materials and Methods and Supplemental Fig. 6A-C, for convenience we symbolized these sites as FoxA3*, ChIP-seq data was obtained from (Iwafuchi-Doi et al. 2016) GSE57559). We found enrichment of Foxa3* binding in the promoters of genes downregulated in the FoxA triple null (Supplemental Fig. 6D-E). We also show a significant association between FoxA3* distal regulatory regions and the downregulated genes using the HiC map of liver chromatin generated by Kim and colleagues (Kim et al. 2018) (GSE104129, Supplemental Fig. 6F-G).

### HNF4α binding is dependent on the presence of FoxA at co-bound sites

Since FoxA proteins play a key role in chromatin remodeling during development (Li et al. 2012; Zaret and Mango 2016), we wondered if the FoxA proteins act only as pioneer factors during fetal organogenesis or if they are continuously required for the hepatic gene regulatory network. We focused on the effects of FoxA depletion on HNF4α binding, because HNF4α is known to have key role in hepatic maintenance and since it has a wide binding pattern across most of hepatic enhancers. Indeed, 61% of FoxA binding sites (FoxA3*) are also occupied by HNF4α (Fig. 4A) and 37% of genes down-regulated in the FoxA triple null livers are also HNF4α dependent (Fig. 4B and Supplemental Fig. 7A). In order to examine the effect of FoxA depletion in the adult liver on HNF4α binding, we performed ChIP-seq for HNF4α on FoxA triple null and control livers. We note that HNF4α protein levels are maintained in FoxA triple null livers, excluding a trivial explanation for loss of HNF4α binding (Supplemental Fig. 7B). We found HNF4α occupancy at binding sites not shared with FoxA binding sites to be similar between mutants and controls, as expected (Fig. 4C). However, at FoxA sites (FoxA3*) co-bound by HNF4α, HNF4α occupancy was dramatically reduced at 40.6% percent of the sites, especially those with relatively weak HNF4α binding (Fig. 4D). Thus, FoxA is required not only during the establishment of the hepatic primordium for the generation of the hepatic enhancer landscape, but also for maintaining the adult hepatic regulatory network by constantly enabling the binding of HNF4α.

**Figure 4.**
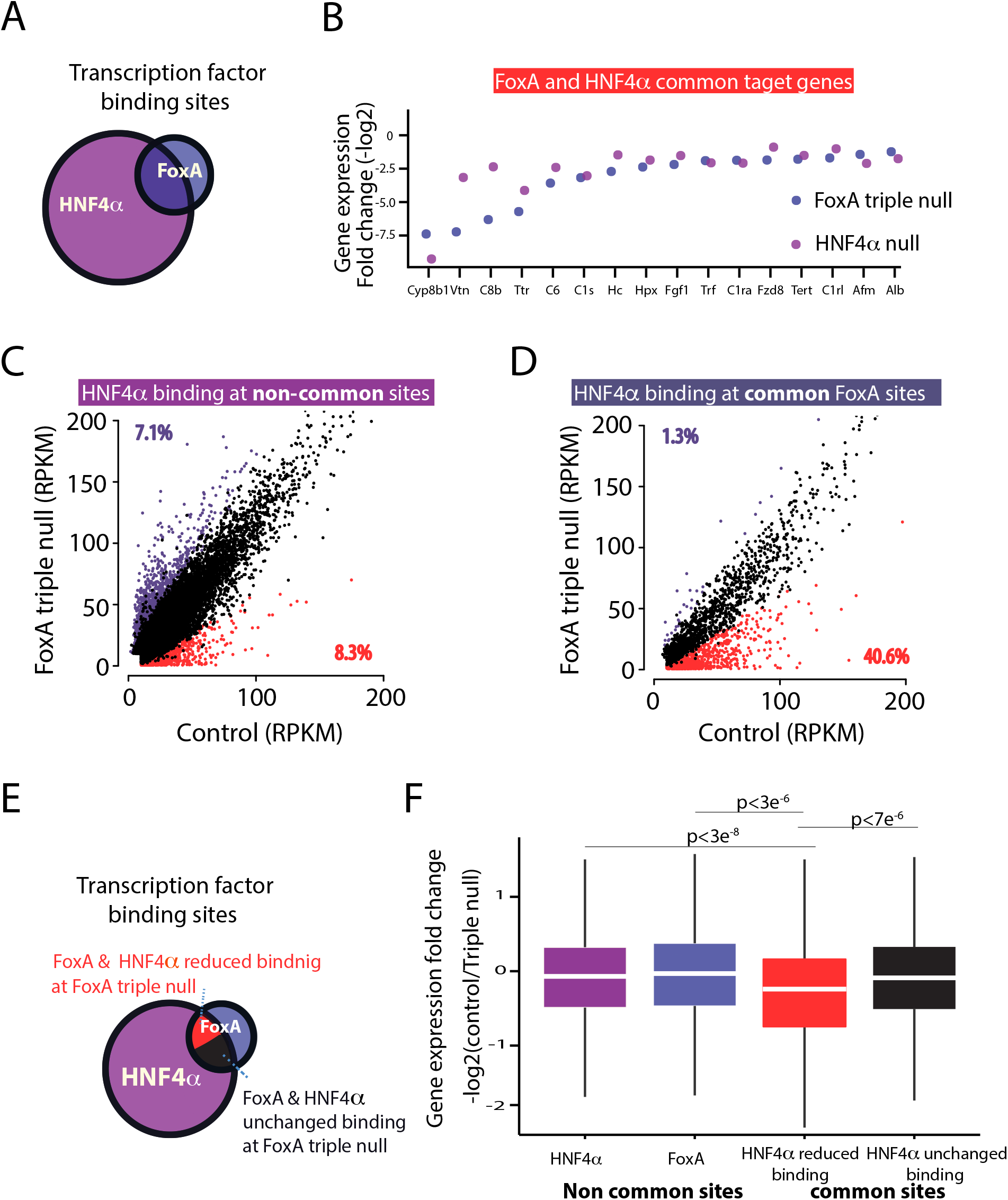
HNF4α binding is dependent on FoxA proteins in co-bound sites. A. Venn diagram of FoxA (FoxA3*) and HNF4α binding sites in the adult liver. B. Transcript levels of key liver genes down-regulated in the FoxA triple null or in adult livers depleted for HNF4α. Presented is fold change of mutants compared with controls for both models. C. Scatter plot of HNF4α binding values (RPKM, reads per kb per million) of FoxA triple nulls compared to controls in HNF4α sites that do not overlap with FoxA (n=4 for controls or mutants). D. Scatter plot of HNF4α binding values of sites with FoxA occupancy. Increased or decreased binding is purple or red, respectively. FDR< 0.05. Each group is the mean of 4 samples. E. Venn diagram showing the proportion of 4 groups of binding sites: non-common sites of either HNF4α or FoxA proteins, common sites that show no loss of HNF4α occupancy, and common sites with loss of HNF4α binding. F. Box plot demonstrating fold change of the nearest gene in the FoxA triple nulls compared with controls for each of the groups indicated in E (Wilcoxon-Mann-Whitney test).

Next, we examined the association between the reduction of HNF4α occupancy at co-bound sites and genes expression. For this purpose, we divided the FoxA and HNF4α binding sites into four classes: non-common sites of either HNF4α or FoxA; common sites that show no loss of HNF4α occupancy; and the common sites with loss of HNF4α binding (Fig. 4E). Strikingly, only common sites with reduced HNF4α binding showed a statistically significant decrease in expression of nearest genes (Fig. 4F). This finding shows that FoxA generates its strongest effects on gene expression in the adult liver at enhancers also bound by HNF4α, in accordance with the view that enhancers are usually occupied by clusters of transcription factors (Spitz and Furlong 2012).

### FoxA maintains enhancer activity, nucleosome positioning, and chromatin accessibility at associated enhancers

Next, we performed ChIP-seq on H3K27ac and H3K4me1, the two main chromatin markers associated with enhancer activity, on the livers of FoxA triple nulls and controls. We focused on HNF4α and FoxA co-bound sites, divided into those with loss of or unchanged HNF4α binding in the FoxA triple nulls. Strikingly, we found that 69.5% of FoxA/HNF4α co-bound regions with loss of HNF4α exhibit a dramatically reduced H3K27ac signal, while this was the case for only 8% of co-bound sites with unchanged HNF4α occupancy (Fig. 5A-B and Supplemental Fig. 8A, FDR < 0.05). In addition, we showed that 20.1% of the co-bound sites with loss of HNF4α binding have a reduced signal for another important enhancer mark, the H3K4me1 modification (Fig. 5A-B and Supplemental Fig. 8A, FDR < 0.05). Strikingly, the H3K4me1 mark, clearly present on two nucleosomes flanking the transcription factor binding sites in the control liver as typically for primed or active enhancers (Fig. 5A), collapsed onto a single nucleosome in the vast majority of the FoxA/HNF4α co-bound enhancers (Fig. 5A) thus showing a shift in nucleosome positioning. However, of the co-bound sites with unchanged HNF4α binding, only 1.9% displayed a reduced H3K4me1 signal, and no change in nucleosome positioning (Fig. 5A-B and Supplemental Fig. 8A, FDR<0.05). Next, we performed ATAC-seq in order to address the mechanism by which FoxA maintains enhancer activity. We found reduced accessibility at 34.4% of FoxA sites co-bound by HNF4α (Fig. 5C, FDR <0.05), while only 0.4% of sites with no decrease in HNF4α binding displayed reduced accessibility (Supplemental Fig. 8B). Thus, the FoxA proteins are continuously required to maintain nucleosome positioning, chromatin accessibility, and active enhancer marks at key regulatory sequences that are required for liver viability, establishing the FoxA factors as active enhancer guards.

**Figure 5.**
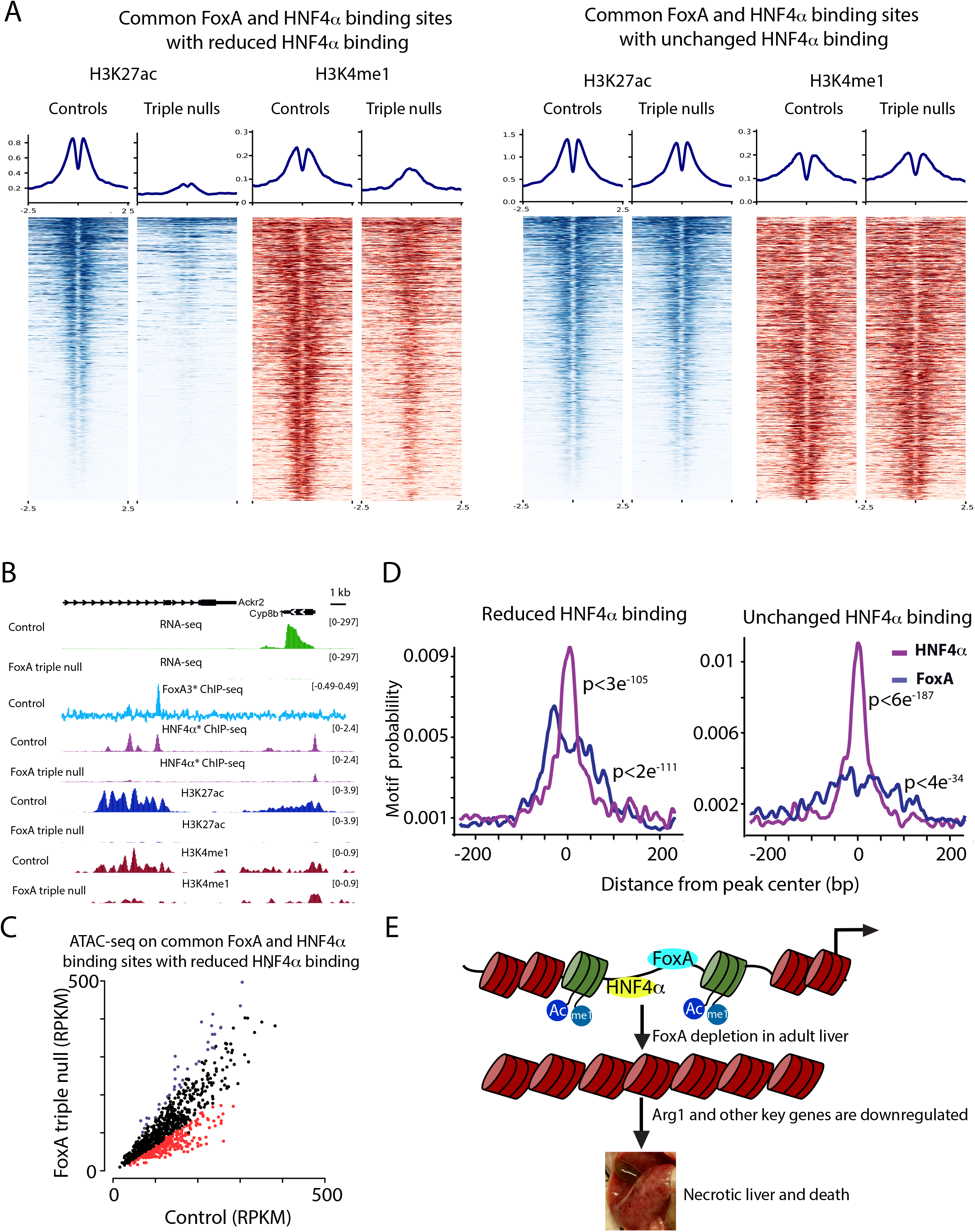
FoxA is required to maintain active enhancer, nucleosome positioning and chromatin accessibility at co-bound HNF4α/FoxA binding sites. A. Heatmap and quantification of H3K27ac and H3K4me1 signal in HNF4α/FoxA common binding sites that show no loss of HNF4α occupancy, and the common sites with loss of HNF4α binding (n=2, for controls or mutants). B. HNF4α/FoxA common site next to the Cyp8b1 gene in which loss of FoxA binding results in loss of expression, HNF4α binding and altered H3K27ac and H4K4me1 signals. C. Scatter plot of ATAC-seq RPKM values in FoxA triple null livers compared to controls in co-bound HNF4α/FoxA sites that show loss of HNF4αbinding. Decreased accessibility sites are indicated as red dots. FDR< 0.05 (n=3 for controls or mutants). D. Motif frequency as a function of distance from peak center (bp) of the binding sites indicated in A using Centrimo. E. Model for the role of FoxA proteins in maintaining enhancer activity, chromatin accessibility, and gene expression in the adult liver.

Lastly, since we observed a strong effect of FoxA on H3K27 acetylation and HNF4α binding at certain sites but not in others, we performed motif frequency analysis using Centrimo, as we hypothesized that binding site density could be a determining factor differentiating between enhancers that are sensitive to FoxA loss versus those that are resistant. Indeed, we found that the stable FoxA/HNF4α enhancers showed much higher frequency for the HNF4α motif over the FoxA motif (Fig. 5D), while those that had lost HNF4α binding and enhancer marks displayed equal frequencies of the two binding elements.

## Discussion

In conclusion, we discovered a set of critical metabolic pathways and genes that are fully dependent on FoxA proteins in terminally differentiated hepatocytes and that are necessary for survival. Many of these key genes were never before described as FoxA targets. Among these are *Arg1*, plasminogen, and *Apoa1*. Our data clearly show that the FoxA proteins are not exclusively acting as competency factors required only during liver development, but rather continuously maintaining critical enhancer function. We have also established that the hepatic gene regulatory network is not resilient to FoxA loss. We suggest a model in which FoxA proteins are continuously required to maintain enhancer activity and nucleosome positioning at enhancers with a high density of FoxA motifs. This binding enables occupancy by other transcription factors and without FoxA proteins many critical established enhancers collapse, leading to near complete loss of gene activity (Fig. 5E).

FoxA is known to play a key role in the development of many organs including pancreas, intestine and lung (Wan et al. 2005; Gao et al. 2008; Ye and Kaestner 2009). It is possible that in each of these organs removal of the FoxA proteins will result in lethality and organ degradation. If correct, this paper demonstrates a phenomenon that is applicable for many biological systems and could identify additional metabolic pathways and gene networks. Moreover, in the current paper, we investigated the role of FoxA in normal homeostasis using complete deletion. Future experiments using the same approach on pathological conditions such as cancer, metastasis and fatty liver have the potential to find new roles of FoxA proteins in pathogenesis.

Our finding that the paradigm pioneer factor is continuously required to maintain chromatin accessibility in the mature organ, suggests that this is the case for other mammalian pioneer factors. This is unlike Zelda that was shown in the fly to be restricted to enhancer activation at early developmental stages. It is to be determined if true pioneer factors such as Zelda exist in mammals, or of the settler model applies to more mammalian transcription factors.

## Materials and methods

### Animals

All animal procedures were approved by the Institutional Animal Care and Use Committee (IACUC) at the University of Pennsylvania. The derivation of the *FoxA1*^*loxP/loxP*^; *FoxA2*^*loxP/loxP*^; *FoxA3*^*−/−*^ as well as HNF4α mice has been reported previously (Kaestner et al. 1998; Sund et al. 2000; Parviz et al. 2002; Gao et al. 2008). Male mice at ages 8-12 weeks were Jugular vein injected with *10*^*11*^ particles of AAV expressing either Cre recombinase (Addgene 107787-AAV8) or GFP (Addgene 105535-AAV8) under the control of the hepatocyte-specific thyroid-binding globulin (Tbg) promoter. Mice were genotyped by PCR of tail DNA as described in the above-mentioned papers. Validation of HNF4α depletion was shown by Armour and colleagues (Armour et al. 2017).

### Metabolite Measurements

Either plasma samples and/or a neutralized perchloric acid (PCA) extract prepared from liver were used for metabolites measurement. The concentration of amino acids was determined by Agilent 1260 HPLC system, using precolumn derivatization with *o*-phthalaldehyde (Nissim et al. 2012; Nissim et al. 2014). Organic acids levels were determined by the isotope-dilution approach and GC-MS system (Weinberg et al. 2000). Carnitine, Acyl-Carnitine, and b-OH-butyrate levels were determined by Agilent LC/MS 6410 Triple Quad system as described (Nissim et al. 2012). Total plasma urea level as in (Nissim et al. 2014) and protein level in liver extract as in (Nissim et al. 2012; Nissim et al. 2014).

### Western Blot Analysis

About 30 mg of frozen liver were added to 500 μL of RIPA Buffer (150 mM NaCl, 50 mM Tris-HCl pH 8.0, 1% Triton X-100, 1 mM PMSF, 0.5% Sodium Deoxycholate, 0.1% SDS, 1X cOmplete protease inhibitor cocktail). Tissue was homogenized in a Tissue Lyser II (Qiagen) using 5 mm stainless steel bead (Qiagen) at 20 Hz for 4 minutes. Following homogenization, the samples were sonicated using a Standard Biruptor (diagenode UCD-200) with the following parameters: Intensity = High (H), Multitimer = OFF: 30 sec, ON: 30 sec, Total timer = 2.5 minutes. Next, the samples were incubated at 4°C for 30 minutes with rotation. Following incubation, samples were centrifuged for 15 minutes at 15,000 RCF and the supernatant was transferred to a new tube. Then, protein quantification using BCA Protein Assay Kit (Thermo Fisher) was done. 50 μg of protein was brought up to 6.5 μL with DDW and added to 2.5 μL of 4X LDS Sample Buffer (Thermo Fisher) and 1 μL of NUPAGE reducing agent (Thermo Fisher). Samples were heated at 70°C for 10 minutes for denaturation and loaded onto a precast 4-12% mini bis-tris gel (Thermo Fisher) using the XCell SureLock Mini-Cell system (Thermo Fisher). Samples were run with 1X MOPS SDS running buffer (Thermo Fisher) with 500 μL of antioxidant (Thermo Fisher). The gel was run at a constant 200 V for 50 minutes. Proteins were transferred to a PVDF membrane (Bio-rad) using a semi-wet method. 1 L of transfer buffer was prepared with 50 mL of 20X NUPAGE transfer buffer (Thermo Fisher),100 mL of methanol, 1 mL of antioxidant, and 849 mL of H_2_O. Gels were transferred using XCell SureLock Mini-Cell System at 30 V for 1 hour. Following the transfer, the membrane was washed with TBST (1X TBS (BioRad), 0.1% Tween-20) for 1 hour. The membrane was blocked with 5% nonfat dry milk (Labscientific) in TBST for 1 hour. Following blocking, membrane was washed in TBST for 30 minutes. The membrane was incubated with the primary antibody diluted at a ratio of 1:1000 in 5% milk in TBST overnight at 4°C with the following antibodies, anti-FOXA1 (Abcam ab23738), anti-FoxA2/HNF3β (D56D6) XP® Rabbit mAb (Cell Signaling Technology #8186) and anti HNF4a (Abcam ab181604). The following day, the membrane was washed in TBST for 30 minutes and then incubated with secondary antibody diluted at a ratio of 1:5000 in 5% nonfat dry milk in TBST for 1 hour at room temperature. Then it was washed in TBST for 30 minutes. Protein signal was measured by chemiluminescence with the application of West Dura Extended Duration Substrate (Thermo Fisher). Following development, the membrane was washed in TBST for 30 minutes, changing the TBST and stripped in Western Blot Stripping Buffer (Thermo Fisher) at room temperature for 45 minutes. Membrane was reprobed following the protocol’s previous detail using an anti-α Tubulin antibody (Santa Cruz Biotechnologies sc-23948).

### Immuno-histochemistry

Paraformaldehyde fixed liver tissues were paraffin-embedded and sectioned. To deparaffinized tissue, slides were incubated for 15 minutes at 56°C and then submerged in xylene for 5 minutes twice. Tissue sections were rehydrated and then slides were quickly dipped in H_2_O and then incubated in PBS for 5 minutes. To retrieve antigens, slides were submerged in 1X R-Buffer A (Electron Microscopy Sciences) and loaded into a 2100 Antigen Retriever (Proteogenix). After reaching the target temperature, slides were allowed to cool for 2 hours. Slides were rinsed with running water for 5 minutes and then incubated in PBST (1X PBS, 0.1% Tween-20) for 5 minutes. Slides were then incubated in PBS for 5 minutes three times. Slides were submerged in 3% H_2_O_2_ for 15 minutes to quench peroxidases, rinsed with gentle running H_2_O for 5 minutes, and placed in PBS. To block the tissue, Avidin D blocking reagent (Vector Laboratories) was applied to the slide for 15 minutes at room temperature. Following the incubation, slides were placed in PBS. Biotin blocking reagent (Vector Laboratories) was applied to the tissue and slides were again incubated for 15 minutes at room temperature and then washed again in PBS. CAS-Block (Thermo Fisher) was applied to the tissue and the slides were incubated for 10 minutes at room temperature. Ki-67 antibody (ThermoFisher RM-9106) was diluted in CAS Block and applied to tissue and the slides were incubated overnight at 4°C in a humidity chamber. The following day, slides were incubated in PBST for 5 minutes and then in PBS for 5 minutes twice. Secondary antibody was diluted at a ratio of 1:200 in CAS Block and applied to the tissue for 30 min at 37C and then washed twice in PBS for 5 minutes. ABC HRP Reagent (Vector Laboratories PK-7100) was applied to tissue and slides were incubated for 30 minutes in a humidity chamber at 37°C. Slides were washed in PBS twice for 5 minutes. Using the DAB Substrate Kit for Peroxidase (Vector Laboratories), the substrate working solution was applied to tissue and signal development was monitored under a microscope. Slides were rinsed with running water for 10 minutes to stop the reaction. To counterstain tissue, slides were dipped in Hematoxylin Solution (Sigma GHS216) and then rinsed with running water for 5 minutes. Tissue sections were dehydrated and then mounted using Cytoseal XYL (Thermo Fisher).

### RNA extraction and library preparation

A maximum of 500,000 hepatocytes were homogenized in TRIzol Reagent (Thermo Fisher) and incubated at room temperature. Then, chloroform purification (Millipore Sigma C2432) was taking place followed by ethanol precipitation. Later, RNA was purified using a RNeasy Mini Kit (Qiagen) and eluted with DDW. RNA was quantified using a NanoDrop-1000 (Thermo Fisher) and RNA quality was measured using a BioAnalyzer RNA Nano Chip (Agilent). Libraries were prepared using an Ultra RNA Library Prep Kit for Illumina (New England BioLabs) with 150 ng of input RNA. Libraries were sequenced on the Illumina Hiseq 4000. RNA was extracted from the following genotypes: FoxA1^*loxP/loxP*^ FoxA2^*loxP/loxP*^, FoxA1^*loxP/loxP*^ FoxA2^*loxP/loxP*^ Cre Alfp, FoxA1^*loxP/loxP*^ FoxA2^*loxP/loxP*^ FoxA3^−/−^ injected with AAV GFP, FoxA1^*loxP/loxP*^ FoxA2^*loxP/loxP*^ FoxA3^−/−^ injected with AAV Cre.

### ChIP-seq

About 130 mg of frozen liver was minced in 2 mL of cold 1X DPBS (Thermo Fisher 14080055). The volumes of liver samples were brought up to 10 mL of 1X DPBS and 1% formaldehyde. The samples were incubated at room temperature for 8 minutes with rotation. To quench, reaction was brought to 0.125 M glycine and incubated for 5 minutes at room temperature with rotation. Samples were centrifuged at 2500 RCF for 2 minutes at 4°C and the supernatant was discarded and washed once with 1XDPBS. The pellet was resuspended in 1 mL of cold ChIP cell lysis buffer (10 mM Tris-HCl pH 8.0, 10 mM NaCl, 3mM MgCl_2_, 0.5% IGEPAL CA-630, 1X cOmplete protease inhibitor cocktail) and transferred to a 10 mL tissue grinder on ice. The samples were homogenized with a smooth Teflon pestle 20 times and incubated at 4°C for 5 minutes. Following incubation, the samples were centrifuged for 5 minutes at 17,000 RCF at 4°C to pellet nuclei. The supernatant was discarded, and nuclei were resuspended in 1 mL of cold ChIP nuclear lysis buffer (50 mM Tris-HCl pH 8.0, 5 mM EDTA, 1% SDS, 1X cOmplete protease inhibitor cocktail). The samples were sonicated for two rounds using the Standard Biruptor (diagenode UCD-200) with the following parameters: Intensity = High (H), Multitimer = OFF: 30 sec, ON: 30 sec, Total timer = 7.5 minutes with maintaining 4°C. The Bioruptor was cooled down for 15 minutes between rounds. Following sonication, the samples were centrifuged at 17,000 RCF for 10 minutes at 4°C and the supernatant containing sheared chromatin was recovered.

To determine the amount of material required for precipitation we isolated DNA from sonicated chromatin. DNA was quantified using NanoDrop-1000 (Thermo Fisher) and run on a BioAnalyzer High Sensitivity Chip. Following quantification, 10 μg of sonicated chromatin was added to 1 mL of ChIp dilution buffer (16. mM Tris-HCl pH 8.0, 167 mM NaCl, 0.01% SDS, 1.1% Triton-X 100, 1X cOmplete protease inhibitor cocktail). 3 μg of the following antibodies, H3K4me1(Abcam ab8895), H3K27ac (Active Motif 39133) and HNF4a (Abcam ab181604) were added to the samples and the samples were incubated at 4°C overnight with rotation. 40 μL of recombinant protein G agarose beads (Thermo Fisher) were washed three times with 1 mL of ChIP dilution buffer by resuspending beads in the buffer. Following the washes, the beads were resuspended in 75 μL of ChIP Dilution Buffer and 5 μL of BSA (New England BioLabs) and incubated at 4°C overnight with rotation for blocking. Then chromatin samples were added to the blocked beads and the samples were incubated at 4°C for 1 hour with rotation. Samples were centrifuged at 100 RCF for 30 seconds and the supernatant was discarded. The beads were then washed sequentially with each of the following buffers by resuspending the beads in 1 mL of buffer, incubating beads at room temperature for 5 minutes with rotation, pelleting beads, and discarding supernatant: TSE I (20 mM Tris-HCl pH 8.0, 150 mM NaCl, 2 mM EDTA, 0.1% SDS, 1% Triton X-100), TSE II (20 mM Tris-HCl pH 8.0, 500 mM NaCl, 2 mM EDTA, 0.1% SDS, 1% Triton X-100), ChIP Buffer III (10 mM Tris-HCl pH 8.0, 0.25 M LiCl, 1 mM EDTA, 1% IGEPAL CA-630, 1% Sodium Deoxycholate), TE (10 mM Tris-HCl pH 8.0, 1 mM EDTA). Following the washes, the beads were resuspended in 100 μL of elution buffer (0.1 M NaHCO_3_, 1% SDS) and incubated at room temperature for 15 minutes with rotation. 4 μL of 5M NaCl was added to the supernatant and then samples were incubated at 65°C overnight. Next, DNA was isolated from the samples.

Enrichment of IP samples was measured through qPCR using the following program: 95°C for 3 minutes, for 40 cycles, 95°C for 5 seconds, 60°C for 20 seconds, with the primers indicated below. Then, libraries were prepared using the NEBNext Ultra II DNA Library Prep Kit for Illumina. Libraries were sequenced on the Illumina Hiseq 4000.

HNF1α promoter-
F - GCA CTT GCA AGG CTG AAG TC
R - ATT GGA GCT GGG GAA ATT CT
40S
F - AGC GAG CTG TGC TGA AGT TT
R - AGG CTG CTT GGA TCT GGT TA

### ATAC-seq

About 50 mg of frozen livers were minced in 1 mL of cold swelling buffer (10 mM Tris-HCl pH 7.5, 2mM MgCl_2_, 3mM CaCl_2_). Tissue was homogenized with a Dounce tissue grinder (DWK Life Sciences 357544) using the loose pestle 10 times and was incubated at 4°C for 20 minutes. Following incubation, the sample was homogenized 20 times with the tight pestle. 3 mL of cold swelling buffer was added to the lysate and the sample was passed through a 70 μm cell strainer (Corning). Lysate was centrifuged at 600 RCF for 20 minutes at 4°C. The supernatant was discarded, and the pellet was resuspended in 1.8 mL of swelling buffer and 200 μL of glycerol. While being vortexed, 2 mL of lysis buffer (1X swelling buffer, 10% glycerol, 1% IGEPAL CA-630) was added drop by drop to the lysate and the lysate was incubated for 5 minutes at 4°C. Following incubation, 5 mL of lysis buffer was added to the sample and the sample was centrifuged at 700 RCF for 13 minutes at 4°C. The supernatant was discarded, and the nuclei were resuspended in 2 mL of 1X DPBS (Thermo Fisher 14080055). Nuclei were counted using a hemocytometer and 135,000 nuclei were added to a new 1.5 mL microcentrifuge tube and pelleted by centrifugation at 900 RCF for 10 minutes at 4°C. The supernatant was discarded and nuclei were resuspended in 75 μL of 2X TD Buffer (Illumina), 49.5 μL of 1X DPBS, 7.5 μL Tn5 Transposase (Illumina), 1.5 μL of 10% Tween-20, 1.5 μL of 1% Digitonin, and 15 μL of DDW. Nuclei were incubated for 30 minutes at 37°C. Following incubation, DNA was purified using the MinElute Reaction Cleanup Kit (Qiagen) and eluted in 10 μL of EB. 25 μL of High-Fidelity 2X PCR Master Mix (New England BioLabs), 2.5 uL of indexing primer and 10 μL of DDW were added to the 10 μL of purified DNA and the DNA was amplified five cycles using the following PCR program: 72°C for 5 minutes, 98°C for 30 seconds and then 5 cycles at 98°C for 10 seconds, 63°C for 30 seconds, and 72°C for 1 minute. Next, 5 μL of the partially amplified library were removed and used to determine the appropriate number of additional cycles by qPCR with the following mix: 3.85 μL of DDW, 0.5 μL of Indexed Primer, 0.15 μL of SYBR Green I Nucleic Acid Gel Stain diluted 1:100 (Thermo Fisher) and 5 μL of High-Fidelity 2X PCR Master Mix. We used the following qPCR program: 98°C for 30 seconds, then 20 cycles at 98°C for 10 seconds, 63°C for 30 seconds, and 72°C for 1 minute. Additional cycles were determined as one-third of the plateau. The remaining 45 μL of the partially amplified library were further amplified accordingly. Double-sided size selection was performed using Ampure XP beads (Beckman Coulter) by adding 22.5 μL of beads to library and keeping the supernatant. The supernatant was transferred to another tube and then 58.8 μL of beads were added. The beads were kept and resuspended in 20 μL of DDW, from which the final library was eluted. Libraries were sequenced on the Illumina Hiseq 4000.

## Data analysis

### Reference for FoxA binding sites indicated in the paper

While FoxA1 and FoxA2 are nearly identical in amino acid sequence and in their cistromes, FoxA3 has diverged somewhat during vertebrate evolution, which is reflected by a partially divergent hepatic cistrome (Iwafuchi-Doi et al. 2016). Thus, FoxA3 has both unique binding sites and many others shared with FoxA1/A2. Interestingly, mice lacking only hepatic FoxA1/A2 have a much milder phenotype compared to the triple nulls (Li et al. 2009). We reasoned that FoxA3 must be able to partially compensate for loss of FoxA1/A2. Indeed, the number of sites occupied by FoxA3 doubles in the FoxA1/A2 mutant mice through occupancy of sites normally bound by FoxA1/2 (from 1,998 to 3732), indicative of enhancer switching (for convenience, FoxA3 binding sites occupied in FoxA1/A2 null livers are symbolized as FoxA3*, FoxA3 data from (Zaret and Mango 2016)). Since the FoxA triple nulls ablation is lethal and brings to the downregulation of a set of genes that are specific to this model and are not downregulated in partially depleted FoxA models, it suggests that enhancers still bound by FoxA3 in FoxA1/A2 null livers are critical for hepatic survival and are the most relevant for studying the association between FoxA binding and downregulated genes in the triple null. Indeed, we found positive association between FoxA3* binding sites and the down-regulated genes in the FoxA triple null (as shown in Supplementary S6). Therefore, FoxA sites mentioned in the paper are FoxA3* binding sites which are the most relevant binding sites to the observed phenotype. This is also the reason for the relatively low amounts of FoxA binding sites reported in this paper.

### RNA-seq

RNA-seq reads were mapped to the mouse genome (mm9) using Star version 2.5.2a (Dobin et al. 2013) with the flags: “--readFilesCommand zcat --outSAMtype BAM SortedByCoordinate -- outSAMstrandField intronMotif”. Samtools view version 1.1 was used to remove duplicate with the flag “-bhq 255”.

Cufflinks v2.2.1 (Trapnell et al. 2013) was used for gene expression quantification. We used the following flags:” -max-mle-iterations 500000 --max-bundle-frags 100000000”. The reads were mapped against Gencode vM1. Then we ran DESeq2 package on R in order to estimate differential expression significance. We kept only genes with FPKM >1. Differentially expressed genes were defined as genes with adjusted p-value smaller than 0.01 and fold change greater than 1.5. RNA-seq data of Hepatoblasts and Endoderm were derived from Nicetto D et al. (Nicetto et al. 2019), and newborn and 20 days old hepatocytes data were derived from Reizel Y et al. (Reizel et al. 2018). Heatmaps for either up-or down-regulated genes were done with ggplot2 using heatmap.2. Log2 of all FPKM was calculated and for better visualization Log2FPKM values greater than 9 or smaller than 0 were attributed as 9 or 0, respectively. To analyze gene ontology, a list of either down-or up-regulated genes and their log2 fold change compared with controls was input into Ingenuity Pathway Analysis (Qiagen). Motif frequency analysis was done with CentriMo (Bailey and Machanick 2012). Gene example figures were done with ggplot2 on log2FPKM values. Zonal expression data were derived from (Halpern et al. 2017).

### ChIP-seq

Single-end sequencing reads were trimmed with Cutadapt and aligned to mouse genome (assembly mm9) using Bowtie2 version 2.3.4.1. PCR duplicates were removed using picard.jar tool and the command MarkDuplicates with the following flags:”REMOVE_DUPLICATES=true VALIDATION_STRINGENCY=LENIENT”. Peak calling was done with macs2 version 2.1.1.20160309, peak calling for HNF4α was done on united files of all control samples using default macs2 parameters. In order to compare differential binding between controls and FoxA triple nulls we used DiffBind version 2.10.0 (Ross-Innes et al. 2012). Peaks were defined as ±250 base pairs from peak center. Normalization was done with DBA_SCORE_RPKM_FOLD and DB.DESeq2 was used to find statistically significant differences between groups. We took only binding sites with an RPKM average larger than 10 in controls or FoxA triple null and counted as significant only sites with FDR < 0.05. The same parameters were used for HNF4α binding comparison, H3K27ac, H3K4me1 and ATAC-seq signal.

Bam files were converted to Bedgraph files after normalizing to 1 million reads per library using BEDtools (“genomecov”) and then converted to bigwig format using UCSC toolkit (“bedGraphToBigWig”). For control and FoxA triple null groups, an average bigwig file was also generated from the replicates (“bigWigMerge“). Bigwig files were loaded on the WashU browser for visualization. Deeptools (version 2.5.7, “computeMatrix” and “plotHeatmap”) was used to generate enrichment heatmaps surrounding selected regions. Motif frequency analysis was performed using Centrimo version 1.5.0. (Bailey and Machanick 2012).

### ATAC-seq

Paired-end sequencing reads were aligned to mouse genome (assembly mm9) using Bowtie2 allowing ends to be soft clipped. Only properly paired alignments were retained for downstream analysis. PCR duplicates were removed using SAMtools (“rmdup”). Reads mapped to mitochondria and ENCODE blacklist regions were removed. All reads aligning to the positive strand were offset by +4 bp, and all reads aligning to the negative strand were offset −5 bp. Comparison between controls and FoxA triple null was conducted using DiffBind in a similar way to ChIP-seq processing. Quality control metrics were evaluated before alignment using FastQC (version 0.11.8) and after alignment using QualiMap (version 2.2) to ensure comparable quality of sequencing libraries.

### Nearest genes analysis

To analyze association between FoxA/HNF4α binding sites and downregulated genes in the FoxA triple nulls, we identified nearest gene using Homer AnnoatatePeaks.pl. Then, we filtered out genes with distance greater than 1MB and genes with RPKM values lower than 1. We chose 1MB as a cutoff because HiC analysis showed that at this distance there is still enrichment between FoxA3* binding sites and the promoters of downregulated genes. For each gene −log2 of fold change between FoxA triple null over controls was calculated. Identical amounts of genes were randomly sampled for each group.

### HiC analysis

In order to find association between FoxA binding sites and downregulated genes in the FoxA triple null using HiC maps, we took advantage of liver HiC maps generated by Kim et al (Kim et al. 2018). We computed the signal between FoxA binding sites and TSS of the above-mentioned genes using the following approach. We binned the genome into 5KB tiles, each with a specific interaction scores, measuring interaction with other tiles on the same chromosome. We took all score values that connect tiles overlapping with FoxA3* with tiles of promoters of downregulated or unchanged genes. Statistical test was calculated on all possible interactions.

### Distribution of Materials, Data, and Code

Distribution of Materials: Mouse strains are available from MMRRC.

Deep Sequencing Data: All data have been deposited in the Gene Expression Omnibus (GEO) under accession number GSE140423.

## Acknowledgments

We thank Dr. Mitch Lazar for critical suggestions on the manuscript and the Lazar lab for the RNA-seq data from HNF4α-deficient liver. We thank our lab members for technical help and fruitful discussions. We thank Dr. I. Nissim, Y. Daikhin, O. Horyn, and Dr. Ilana Nissim for performing the analysis of urea, protein, amino and organic acids, carnitine and Acyl-Carnitine levels at the Metabolomics Core Facility at the Children’s Hospital of Philadelphia. We thank Dr. C.A. Assenmacher for pathology analysis of liver sections and providing relevant images. We acknowledge Dr. Jonathan Schug for his advice on bioinformatics analysis and thank Nicholas J. Hand for fruitful discussions.

## Author contributions

YR and KK – Conceptualization and writing. YR, AM, EW – Methodology. YR, AM – Resources. YR,LG,YL,EM,KK-Data curation and visualization. KK and AW-Supervision. KK – Funding acquisition;

