## Supplemental Figures for "Collapse of the hepatic gene regulatory network in the absence of FoxA factors"

**Supplemental Information**

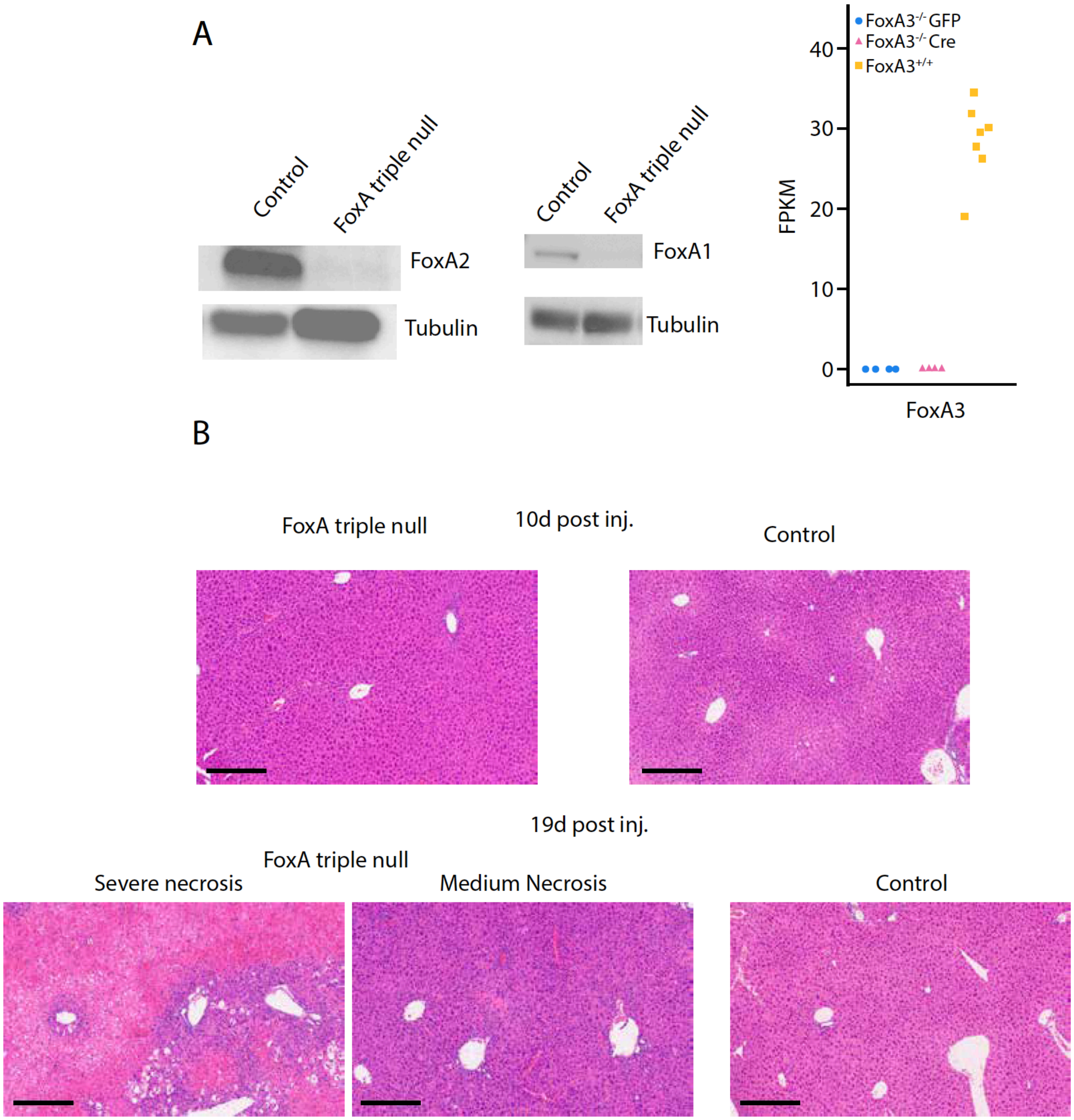


**Supplemental Figure 1. FoxA triple null validation and liver morphology.** A. Western blots on FoxA1 and FoxA2 and mRNA expression levels of FoxA3 in the triple null compared with controls. B. Hematoxylin and Eosin staining for FoxA triple nulls and controls, 10 and 19 days after Cre or GFP induction. Scale is 200 µm.

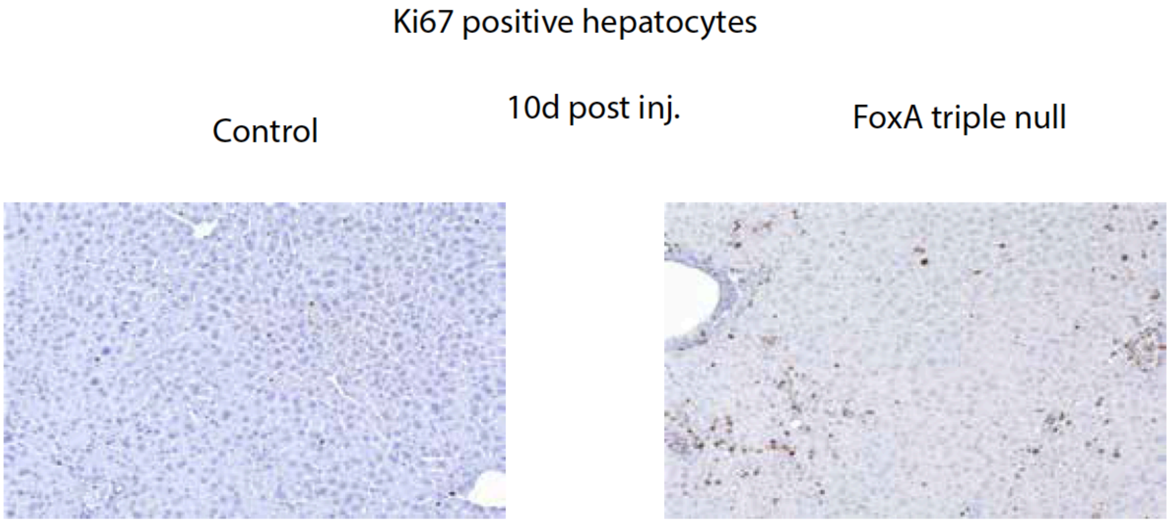


**Supplemental Figure 2. Higher proliferation rates in FoxA triple nulls.** Immunohistochemistry for Ki67 in FoxA triple nulls and controls, 10 days after Cre induction.

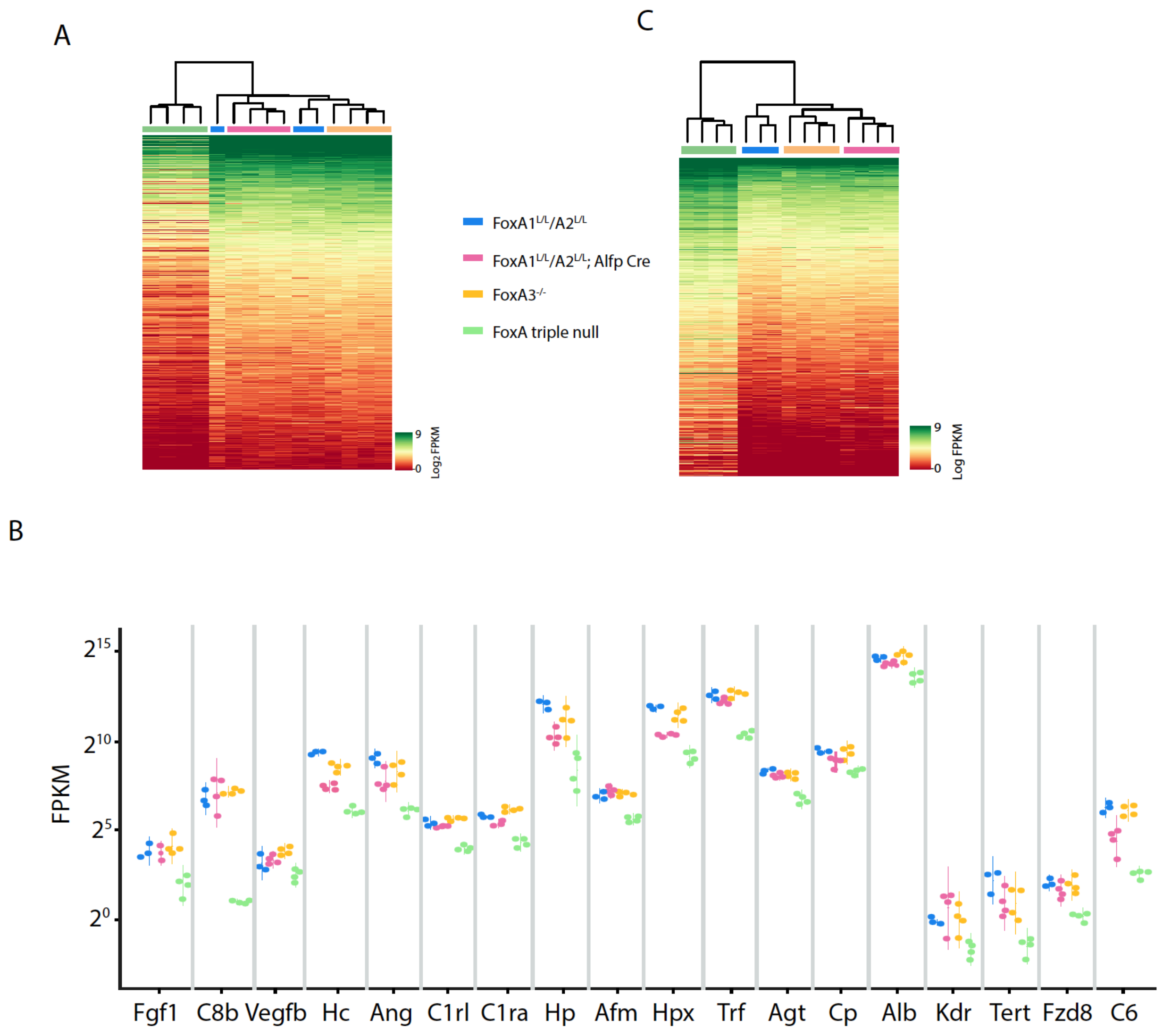


**Supplemental Figure 3. Gene expression profile of FoxA triple nulls.** A. Heatmap of 663 down regulated in FoxA triple null livers seven days after Cre induction compared to the partially FoxA depleted models indicated (adjusted p <0.05, fold change > 1.5). B. Gene expression levels of key liver genes downregulated in the FoxA triple nulls compared to partially depleted FoxA models (adjusted p value < 0.05, fold change > 1.5). C. Heatmap of 513 induced genes in FoxA triple nulls compared with partially depleted FoxA models (adjusted p value < 0.05, fold change > 1.5).

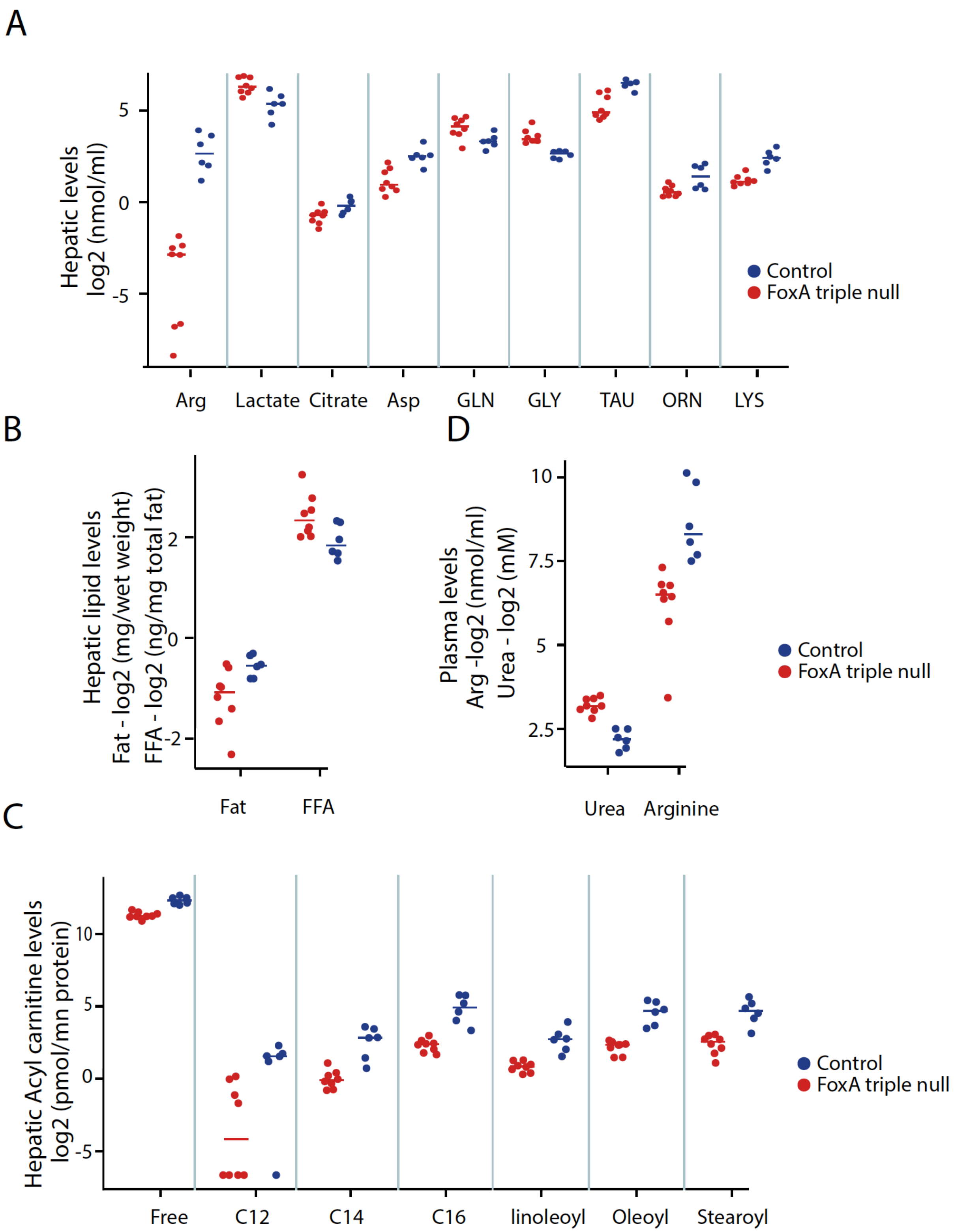
A
Hepatic levels
log2 (nmol/ml)
5
0
-5
Arg
Lactate
Citrate
Asp
GLN
GLY
TAU
ORN
LYS
Control
FoxA triple null
B
Hepatic lipid levels
Fat - log2 (mg/wet weight)
FFA - log2 (ng/mg total fat)
2
0
-2
Fat
FFA
D
Plasma levels
Arg -log2 (nmol/ml)
Urea - log2 (mM)
10
7.5
5
2.5
Urea
Arginine
Control
FoxA triple null
C
Hepatic Acyl carnitine levels
log2 (pmol/mn protein)
10
5
0
-5
Free
C12
C14
C16
linoleoyl
Oleoyl
Stearoyl
Control
FoxA triple null

**Supplemental Figure 4. Metabolomics profiling of FoxA triple null.** FoxA triple null metabolomics profiling of hepatic amino acids, lipids, and Acyl carnitine (A-C) as well as plasma levels of Urea and Arginine (D). $p < 0.05$ Mann-Whitney-Wilcox test for all comparisons presented.

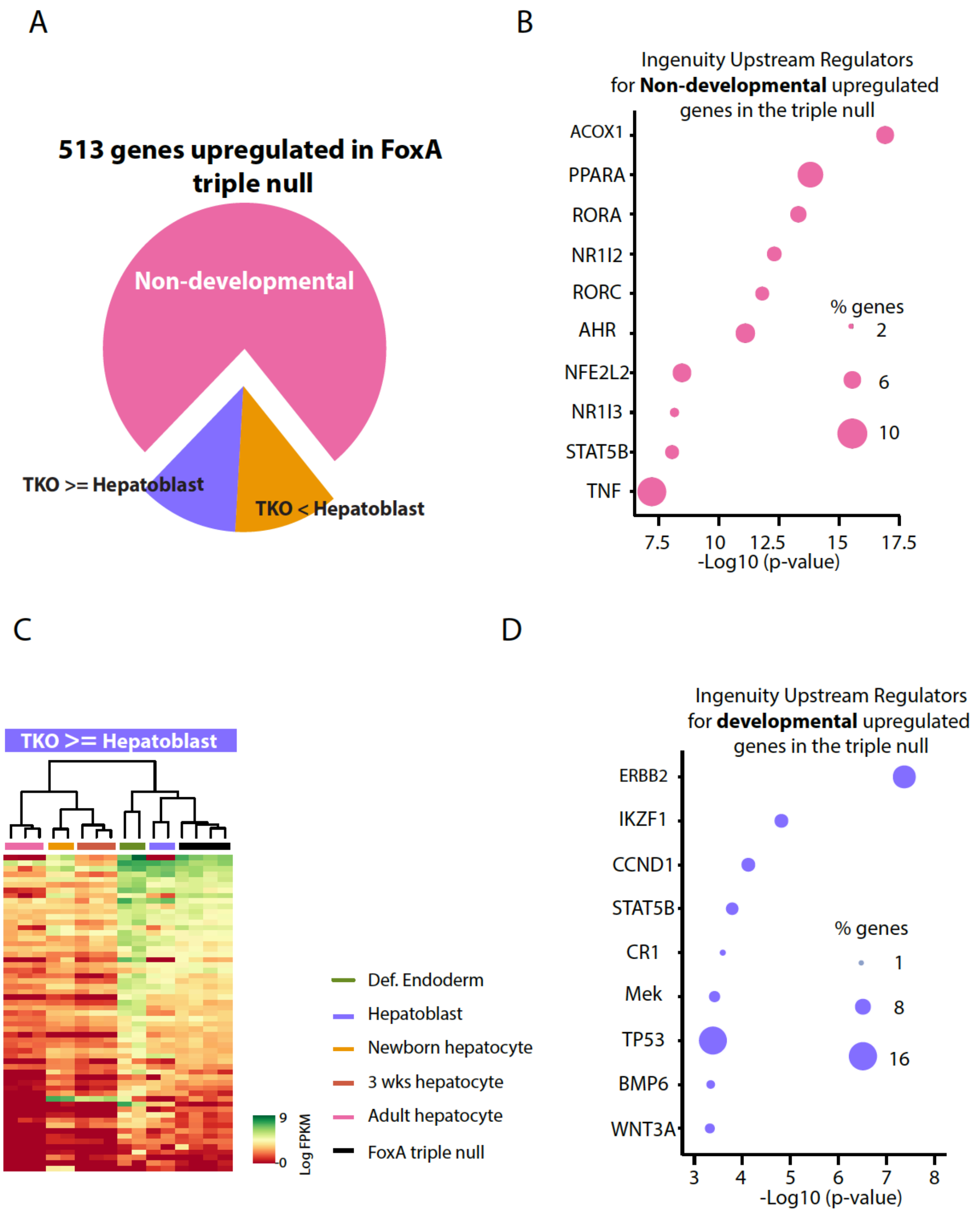


**Supplemental Figure 5. Adult ablation of FoxA factors upregulates genes associated mainly with stress response and metabolism.** A. Only a minority of the upregulated genes in the triple null are repressed during liver development, and an even smaller portion are reactivated in the

FoxA triple nulls back to total hepatoblast levels. B. Heatmap of the few upregulated genes whose expression is elevated back to prehepatic stage. C-D. Upstream regulators of developmental and non-developmental upregulated genes in the FoxA triple nulls. Non-developmental genes are mainly associated with upstream regulators associated with metabolism and stress response, and developmental genes are mainly associated with proliferation.

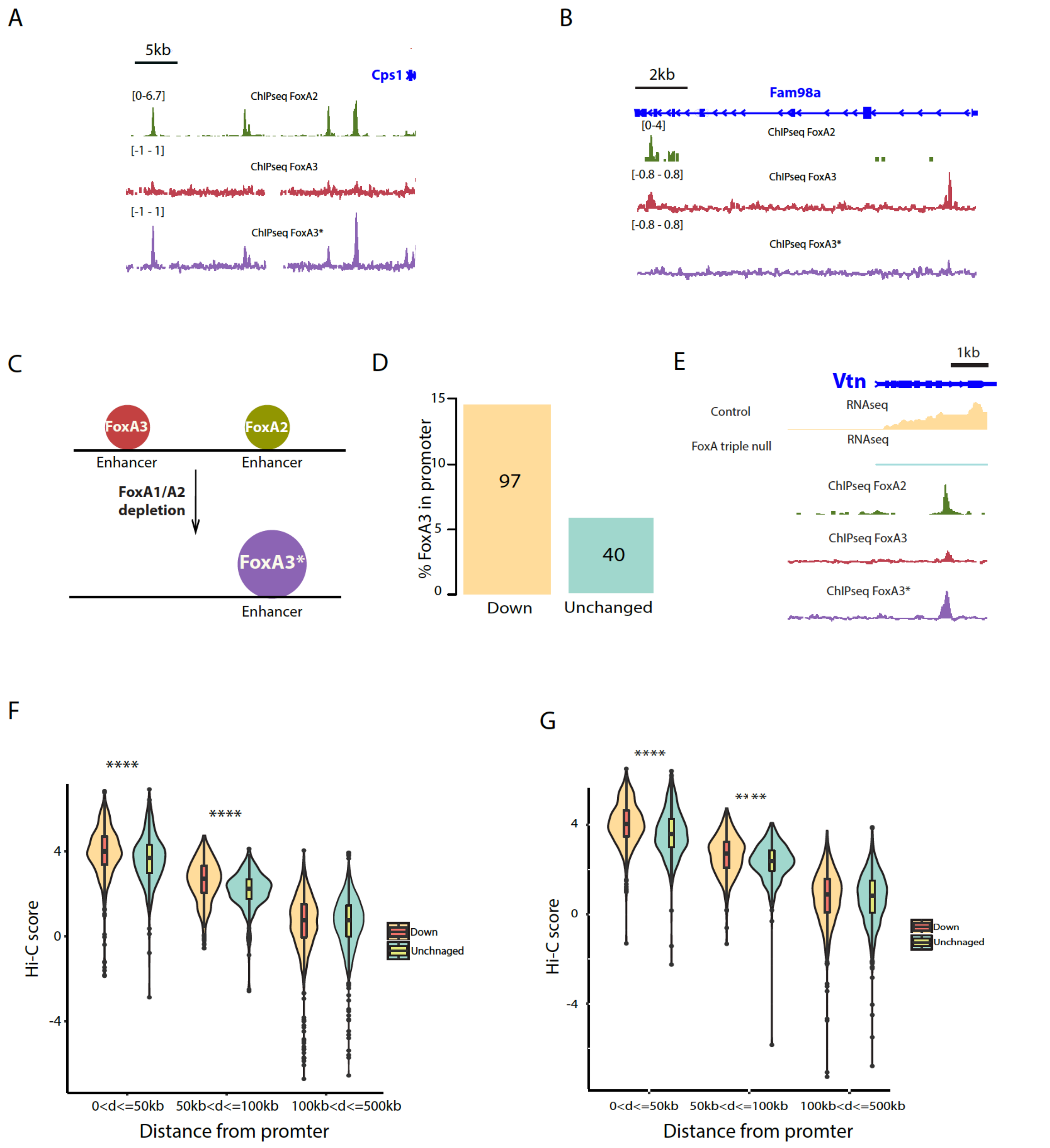


**Supplemental Figure 6. FoxA3* binding sites are associated with down-regulated genes in FoxA triple null.** A-B. Two examples of enhancer switching of FoxA3 in FoxA1/A2 double mutants. C. A model showing that FoxA3 doubles its hepatic binding sites in FoxA1/A2 double mutants. Most of these sites overlap with FoxA1/A2 binding sites. D. Percentage of FoxA3*

binding promoters in downregulated genes in FoxA triple null compared with controls and genes that are unchanged in FoxA triple null ($p < 0.0001$, proportional test, numbers in bars represent number of genes). E. An example of FoxA3* binding next to Vtn promoter. F-G. Violin plots showing higher HiC contacts between FoxA3* binding sites and promoters of downregulated genes in FoxA triple null compared with promoters of unchanged genes. (**** means $p<0.00001$ one sided Wilcoxon test, shown are two different HiC data sets taken at different circadian time points).

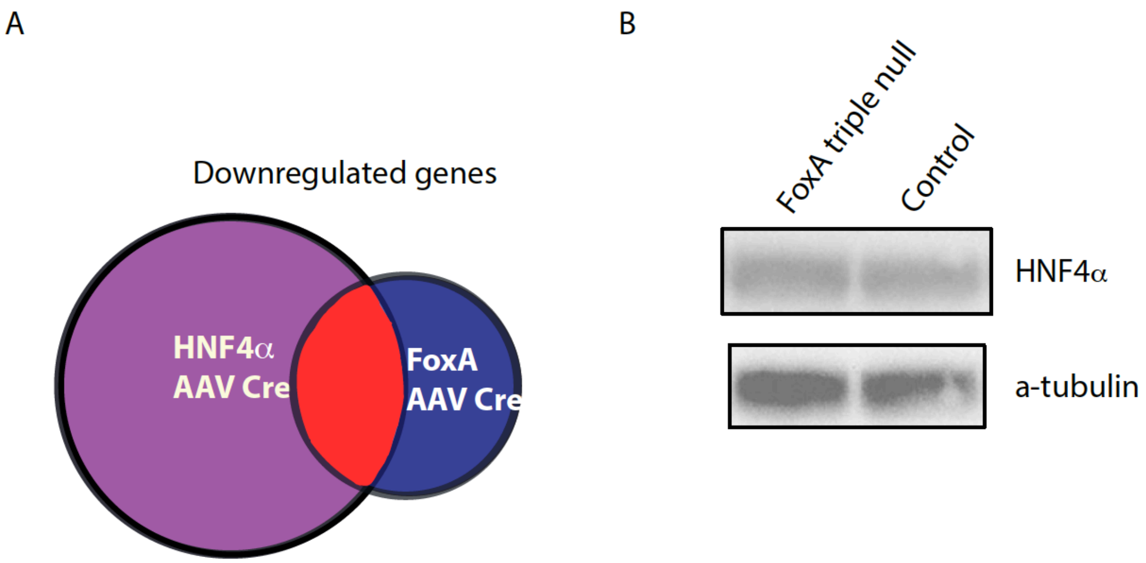


**Supplemental Figure 7. FoxA triple nulls have no effect on HNF4α protein levels, and many genes are coregulated by FoxA and HNF4α**. A. 37% of FoxA triple null repressed genes are also downregulated in the HNF4α deficient liver. B. Western blot showing no change in a HNF4α protein levels in FoxA triple null.

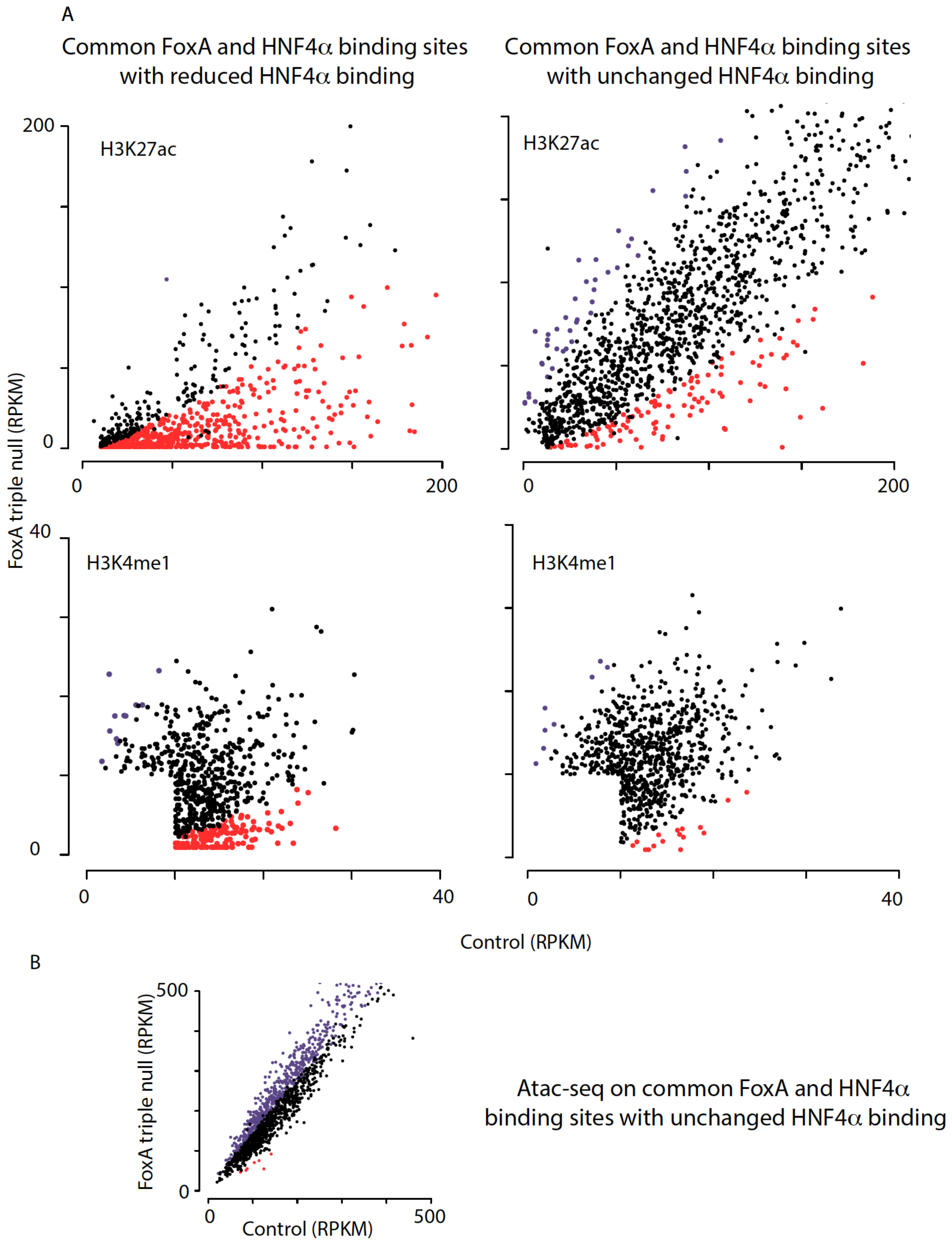


**Supplemental Figure 8. Co-bound FoxA/HNF4α sites loose enhancer markers in the FoxA**

**triple null.** A. Quantification of H3K27ac and H3K4me1 signal in HNF4α/FoxA3* common sites that show either no loss of HNF4α occupancy or loss of HNF4α binding. Increased or decreased H3K27ac or H3K4me1 is marked in purple or red, respectively. FDR< 0.05. Each group contains 2 samples. B. Scatter plot of ATAC-seq RPKM values in FoxA triple null livers compared to controls in co-bound HNF4α/FoxA sites that show no loss of HNF4α binding. Decreased accessibility sites are indicated as red dots and increased with purple. FDR< 0.05. Each group contains 3 samples.
